# Acoustomechanically activatable liposomes for ultrasonic drug uncaging

**DOI:** 10.1101/2023.10.23.563690

**Authors:** Mahaveer P. Purohit, Kanchan Sinha Roy, Yun Xiang, Brenda J. Yu, Matine M. Azadian, Gabriella Muwanga, Alex R. Hart, Ali K. Taoube, Diego Gomez Lopez, Raag D. Airan

## Abstract

Ultrasound-activatable drug-loaded nanocarriers enable noninvasive and spatiotemporally-precise on-demand drug delivery throughout the body. However, most systems for ultrasonic drug uncaging utilize cavitation or heating as the drug release mechanism and often incorporate relatively exotic excipients into the formulation that together limit the drug-loading potential, stability, and clinical translatability and applicability of these systems. Here we describe an alternate strategy for the design of such systems in which the acoustic impedance and osmolarity of the internal liquid phase of a drug-loaded particle is tuned to maximize ultrasound-induced drug release. No gas phase, cavitation, or medium heating is necessary for the drug release mechanism. Instead, a non-cavitation-based mechanical response to ultrasound mediates the drug release. Importantly, this strategy can be implemented with relatively common pharmaceutical excipients, as we demonstrate here by implementing this mechanism with the inclusion of a few percent sucrose into the internal buffer of a liposome. Further, the ultrasound protocols sufficient for in vivo drug uncaging with this system are achievable with current clinical therapeutic ultrasound systems and with intensities that are within FDA and society guidelines for safe transcranial ultrasound application. Finally, this current implementation of this mechanism should be versatile and effective for the loading and uncaging of any therapeutic that may be loaded into a liposome, as we demonstrate for four different drugs in vitro, and two in vivo. These acoustomechanically activatable liposomes formulated with common pharmaceutical excipients promise a system with high clinical translational potential for ultrasonic drug uncaging of myriad drugs of clinical interest.

**One Sentence Summary:** Incorporating a few percent sucrose into a liposome transforms it into an immediately translatable vehicle for noninvasive, on-demand ultrasound-targeted drug delivery.

## INTRODUCTION

Externally-triggered stimulus-responsive drug delivery systems promise targeted, on-demand drug delivery to regions of interest in the body. Ultrasound-gated drug delivery systems are of particular interest as they can leverage recent advances in clinical therapeutic focused ultrasound systems, which can target sonication to millimeter-sized regions of interest throughout the body and are in active clinical use for a variety of applications worldwide (*1–4*). With ultrasonic drug uncaging using such systems, following intravenous infusion of ultrasound-sensitive drug-loaded nanocarriers, focused ultrasound would be applied to a region of interest in the brain or body with a protocol sufficient to induce drug release from the nanocarriers while they are circulating in the blood volume or are resident in the parenchyma of the region of interest (*5–12*). The freed drug would then be able to enter the otherwise unperturbed tissue parenchyma, as it might normally do when given in an unencapsulated form.

Several drug-loaded nanocarriers have been described for enabling ultrasonic drug uncaging including perfluorocarbon-based systems that incorporate drug into the shell of gas-phase microbubbles (*13–18*) or liquid-phase nanodroplets (*13, 19–24*). For microbubble-based carriers, ultrasound-mediated cavitation is the intended mechanism for ultrasound-triggered drug release, which while effective may also induce sonoporation or other forms of injury to the surrounding tissue, raising safety and practical clinical implementation concerns (*14, 15, 25*). For liquid perfluorocarbon nanodroplets, vaporization to a gas phase is many times the proposed drug release mechanism, with cavitation also being involved in instances (*26–28*). For both liquid and gas-based perfluorocarbon systems, since most drugs show negligible solubility in perfluorocarbons, the drug is loaded in the particle shell, limiting the drug loading potential of the system and likely leading to nonspecific release of the agent upon exposure to body temperature plasma via diffusion of the drug across the few nanometers thick particle shell. A non-perfluorocarbon-based ultrasound-triggered delivery system is offered by heat-sensitive liposomes which have a differential melting behavior of some lipid shell components upon heating (*29–32*). In this system, ultrasound or radiofrequency irradiation is applied to heat the medium temperature, inducing drug release from the liposome core. While effective for drug release, heating the tissue to the transition temperature of these liposomes may induce a local heat shock of the background tissue, which is not always desired. Further, maintaining a higher tissue temperature may present challenges in highly vascularized tissue or in organs which move with cardiac or respiratory motion, and necessitates high intensity ultrasound or radiofrequency application, necessitating high performance requirements of the actuating device (*33*). Alternatively, a different liposomal system has been described in which a porphyrin or related molecule is incorporated into the liposome, producing reactive oxygen species upon sonodynamic ultrasound application, inducing leakage of the encapsulated therapeutic (*34*). While this does not necessarily require cavitation or heating, this system does require continuous ultrasound application for several minutes, providing similar technical constraints as with heat-activated liposomes. Another recently-described alternative liposomal system for ultrasound-triggered release includes supercritical carbon dioxide encapsulated into the liposome (*35, 36*), which also raises concern for cavitation related risks and for system stability, as with microbubbles.

With these factors in mind, we aimed to develop a system for ultrasound-triggered drug release that 1) enables high drug loading potential for varied drugs of interest, 2) has minimal drug release without ultrasound, 3) can be activated following intravenous administration and not necessarily intraparenchymal injection, and 4) an ultrasound-activation mechanism that is mechanically-based without necessitating medium heating above body temperature, nor cavitation, and could be activated with a lower duty cycle, shorter ultrasound protocol than is used for heat or sonodynamic activation. As we demonstrate here, we have been able to accomplish this by tuning the acoustic impedance and osmolarity of the liquid core of liposomes to maximize ultrasound responsiveness while maintaining overall stability of the liposome structure without ultrasound. Further, we have been able to formulate these acoustomechanically-activatable liposomes (AAL) with common pharmaceutical excipients while seeing effective ultrasonic drug uncaging with ultrasound intensities within FDA and society guidelines for safe ultrasound application, indicating that this system has high clinical translational potential.

## RESULTS

### Design and development of acoustomechanically activatable liposomes (AALs)

In designing a next-generation ultrasound-activatable drug carrier for ultrasonic drug uncaging, we first considered prior successful formulations that utilize a liquid perfluorocarbon nanodroplet as the ultrasound-responsive element. We noted that in some of these implementations, while cavitation or a liquid-to-gas phase transition may be involved with the drug release mechanism, such cavitation or gas phase induction is not necessarily always present during drug release (*21*). We then hypothesized that we could retain ultrasound sensitivity in the system by replacing the core perfluorocarbon with a liquid droplet that has an acoustic impedance difference with the surrounding medium, to mediate an ultrasound interaction with the droplet relative to the medium. Further, by choosing a replacement liquid that can also be loaded with a drug, we would increase the drug loading potential of the system and likely also the overall stability of drug binding. Additionally, by replacing the perfluorocarbon for more commonly used pharmaceutical excipients, we would increase the clinical translatability of the system. Indeed, several candidate aqueous buffers show significantly varying acoustic impedance measurements in the literature, suggesting that a liposome with an internal compartment composed of a differential acoustic impedance buffer could be optimized for ultrasound response (*37, 38*). Finally, if the internal buffer has a substantially different osmolarity compared to the surrounding medium, this osmotic gradient would induce a fluid shift following the membrane leakiness induced by the initial ultrasound application, thereby accelerating further drug release with ultrasound (Fig. 1A).

**Fig. 1.**
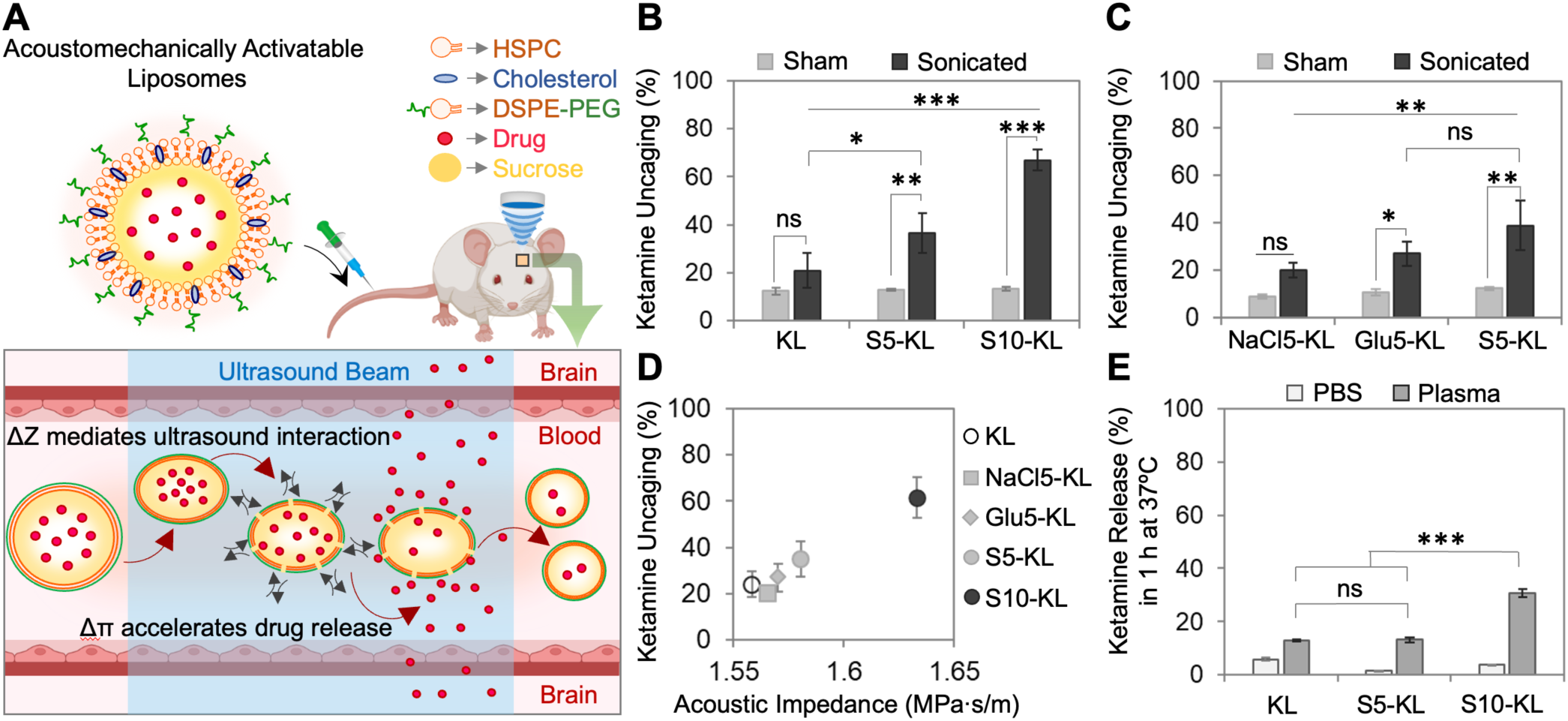
Altering the acoustic impedance of the internal medium of liposomes enhances their ultrasound sensitivity for on-demand drug release. (**A**) Hypothesized scheme of acoustomechanically activatable liposomes (AAL) and proposed mechanism for ultrasonic drug uncaging. Sucrose incorporated into the internal medium of liposomes increases both the acoustic impedance and the osmolarity of the internal buffer. Ultrasound mechanically affects the intravenously injected AAL, due to the difference in the acoustic impedance (ΔZ) of the liposomal internal medium and the surrounding medium, increasing drug permeability across the lipid membrane. Then, the difference in osmotic pressure (Δπ) across the lipid membrane accelerates the rate of water entry into the liposome and drug uncaging into the blood, after which the drug diffuses into the tissue that received sufficient ultrasound intensity. (**B**) In vitro ketamine uncaging from liposomes without (KL) and with 5% (S5-KL) or 10% (S10-KL) added sucrose in the liposome internal medium. Sonicated (250 kHz center frequency, 1.7 MPa peak negative pressure at 25 °C, 25% duty cycle, PRF 5 Hz, 60 s) samples with more sucrose internally release more drug. (**C, D**) In vitro ketamine uncaging from liposomes with internal buffers composed of ammonium sulfate with equiosmolar additional sodium chloride (NaCl5-KL), glucose (Glu5-KL), or sucrose (S5-KL) yields differing amounts of drug release with the same ultrasound protocol, in correlation to their differences in internal acoustic impedances; r^2^=0.97, p = 0.002. (**E**) Spontaneous ketamine release from KL, S5-KL, S10-KL with 1 hr incubation at 37 °C in plasma and buffer shows different stability characteristics, with 10% additional sucrose in the liposome internal medium showing reduced plasma stability at 37 °C. Data presented as mean ± SD of three or more independent experiments. Comparisons between two groups was performed by two-tailed Student’s t-test, and that among multiple groups by one way analysis of variance (ANOVA); ns= non-significant, *p < 0.05, **p < 0.01 and ***p < 0.001.

To implement this hypothetical system, we made liposomes containing varied percentages of sugars in its internal loading battery. We chose to start with liposomes as they are now relatively commonly used drug delivery vehicles formulated with excipients in the FDA Inactive Ingredients Database (*39, 40*). We further chose to incorporate sugars into the internal buffer of the liposome since moderate percentages of sugars have been shown to substantively shift the acoustic impedance of aqueous buffers (*38, 41*).

Additionally, since these sugars are GRAS (Generally Recognized As Safe) excipients for pharmaceutical formulations (*39*), this system would maintain high clinical translational potential for the eventual vehicle. We tested this system with loading of the anesthetic and antidepressant ketamine, a lipophilic amphipathic weak base that we found could be actively loaded into liposomes with an ammonium sulfate loading battery (Table 1). Compared with a ketamine-loaded liposome with no added internal excipient other than the ammonium sulfate loading battery, liposomes incorporating varied percentages of sucrose showed significantly different amounts of ultrasound-induced drug release: with 5% sucrose added to the internal buffer yielding ∼40% drug release with a 250 kHz, 25% duty cycle, 1 min ultrasound protocol, and 10% added sucrose yielding ∼60% drug release with this same protocol, compared to a nonsignificant trend towards ∼20% drug release with no added sucrose (Fig. 1B). Notably, there were no significant differences in key physicochemical properties including size, polydispersity, zeta potential, drug loading, or unencapsulated drug percent (Table 1).

**Table 1.** Typical physicochemical characteristics of different drug-loaded AALs. Note: S5 = 5% additional sucrose, S10 = 10% additional sucrose in the internal buffer of the liposome. The pKa and logP values were derived from PubChem and DrugBank.

| Formulation | Average Size (nm) | PDI | Zeta Potential (mV) | Drug Loading (mg/mL) | Free Drug (%) | Entrapment Efficiency (%) | pKa | LogP |
| --- | --- | --- | --- | --- | --- | --- | --- | --- |
| Ketamine Liposome (KL) |  |  |  |  |  |  | 7.5 | 2.2 |
| KL | 164.07 ± 6.5 | 0.06 ± 0.02 | -45.2 | 3.19 ± 0.17 | 4.96 ± 0.80 | 24-27 |  |  |
| S5-KL | 158.06 ± 5.5 | 0.08 ± 0.01 | -46.3 | 3.41 ± 0.24 | 3.09 ± 1.38 | 25-29 |  |  |
| S10-KL | 160.86 ± 6.0 | 0.06 ± 0.03 | -44.9 | 3.27 ± 0.21 | 3.59 ± 2.02 | 25-28 |  |  |
| NaCl5-KL | 160.36 ± 7.0 | 0.06 ± 0.02 | -48.7 | 3.21 ± 0.14 | 4.62 ± 2.16 | 25-27 |  |  |
| Glu5-KL | 157.86 ± 7.0 | 0.09 ± 0.01 | -47.7 | 3.24 ± 0.20 | 3.24 ± 2.42 | 24-28 |  |  |
| Lidocaine Liposome (LL) |  |  |  |  |  |  | 8.0 | 2.3 |
| LL | 159.90 ± 2.5 | 0.06 ± 0.01 | -49.5 | 4.61 ± 0.07 | 1.82 ± 0.32 | 36-39 |  |  |
| S5-LL | 161.10 ± 2.1 | 0.06 ± 0.02 | -49.1 | 4.50 ± 0.14 | 1.45 ± 0.45 | 36-40 |  |  |
| Bupivacaine Liposome (BL) |  |  |  |  |  |  | 8.1 | 3.4 |
| BL | 165.80 ± 3.2 | 0.09 ± 0.01 | -55.4 | 5.60 ± 0.09 | 3.86 ± 1.02 | 39-42 |  |  |
| S5-BL | 164.38 ± 3.6 | 0.06 ± 0.03 | -49.2 | 5.81 ± 0.12 | 4.02 ± 3.02 | 39-44 |  |  |
| Ropivacaine Liposome (RL) |  |  |  |  |  |  | 8.1 | 2.9 |
| RL | 169.73 ± 3.2 | 0.07 ± 0.02 | -43.3 | 4.32 ± 0.35 | 3.81 ± 2.12 | 38-43 |  |  |
| S5-RL | 166.71 ± 3.8 | 0.08 ± 0.01 | -49.2 | 4.51 ± 0.42 | 4.85 ± 1.07 | 37-41 |  |  |

We next noted that increasing sucrose percentages would yield increases in both the acoustic impedance and the osmolarity of the internal buffer, and the differences in both parameters with the surrounding medium. To help discern which parameter, acoustic impedance or osmolarity, was more contributory to our results we next prepared liposomes with equiosmolar internal buffers, but differential acoustic impedances. Drug-loaded liposomes with internal buffers composed of ammonium sulfate with equiosmolar additional sodium chloride, glucose, or sucrose were prepared and found to have similar physicochemical characteristics with each other (Table 1). Indeed, despite similar internal osmolarities, they showed differing amounts of drug release with the same ultrasound protocol, in correlation to their differences in internal acoustic impedance (Fig. 1B,C). This indicates that acoustic impedance differences across the shell of a liposome likely contribute to the ultrasound-responsiveness of the drug-loaded nanocarrier. Potentially, an internal-external osmolarity gradient then further contributes to increase this drug release following the initial ultrasound-induced membrane leakiness. Notably, while 10% additional sucrose internally significantly increased the amount of ultrasound-induced drug release relative to 5% additional sucrose, it did so at the expense of displaying a significant rate of drug release without ultrasound when incubated with plasma at 37 °C, indicating that there is a relative tradeoff of ultrasound responsiveness and formulation stability with this scheme (Fig. 1D). For further experiments, 5% sucrose added internally was chosen as a relative optimum of ultrasound responsiveness with stability of the formulation with body temperature incubation in plasma.

### Mechanism of AAL-mediated ultrasonic drug uncaging

To further explore the mechanism of drug release with this system, the ultrasound frequency was varied between either 250 kHz or 650 kHz, usual frequencies for transcranial therapeutic ultrasound application clinically (*42, 43*). Significant drug release was seen with 250 kHz sonication, with increasing peak pressure/mechanical index correlating to increased drug release, similar to prior results with liquid perfluorocarbon based systems (Fig. 2A) (*12, 21*). Intuitively, there was increased drug release seen with a higher medium temperature of 37 °C versus 25 °C (Fig. 2A). In contrast, with 650 kHz sonication, only marginal drug release was seen with ultrasound, with a minimal dose-response relationship with ultrasound peak pressure/mechanical index, and with only minimal variation with medium temperature (Fig. 2B). This is in distinct contrast to perfluorocarbon-based systems that show significant drug release across a range of frequencies up to and beyond 1 MHz (*7, 9, 21, 26*). This increased responsiveness at 250 kHz compared to 650 kHz argues against both a thermal or radiation force type mechanism for the drug release, since both thermal and radiation force transfer mechanisms tend to increase with increasing acoustic attenuation coefficients, which in turn increase with the acoustic frequency (*44*). Arguing against cavitation as an underlying drug release mechanism, the liquid phase of the formulation particles has a low vapor pressure and we observe no evidence of wide-band or ultraharmonic acoustic scatter with sonication of the liposomal formulation, in contrast to sonication of gas-phase microbubbles, which readily generate these signals of cavitation (Fig. 2C). By electron microscopy, the formulation particles show the expected spherical morphology, but with no internal precipitation of the drug that is shown to correlate to stability of drug loading with other liposomal formulations. With sonication, the size distribution of the particles decreased, cohering with an ultrasound-induced leak of the internal contents (Fig. 2D). Nonetheless, this formulation shows stability for both drug loading and ultrasound-induced release across months of time in refrigerated storage (Fig. 2E). Together, this frequency response difference and these other characterizations suggest a non-cavitation-based acoustomechanical type drug release mechanism, such as an ultrasound-induced rarefaction or oscillation of the internal droplet relative to the surrounding medium, as mediated by the differential acoustic impedance of the internal droplet, contributing to increased permeability of the liposome membrane to the loaded drug with ultrasound. Notably, effective ultrasound-induced release was seen at body temperature with 250 kHz ultrasound and an in situ peak negative pressure of 0.9 MPa, or a mechanical index of 1.8, which is within FDA and AIUM guidelines for safe ultrasound application (without microbubbles present, as in this application) (*45*). Accordingly, we chose this ultrasound protocol for subsequent evaluations.

**Fig. 2.**
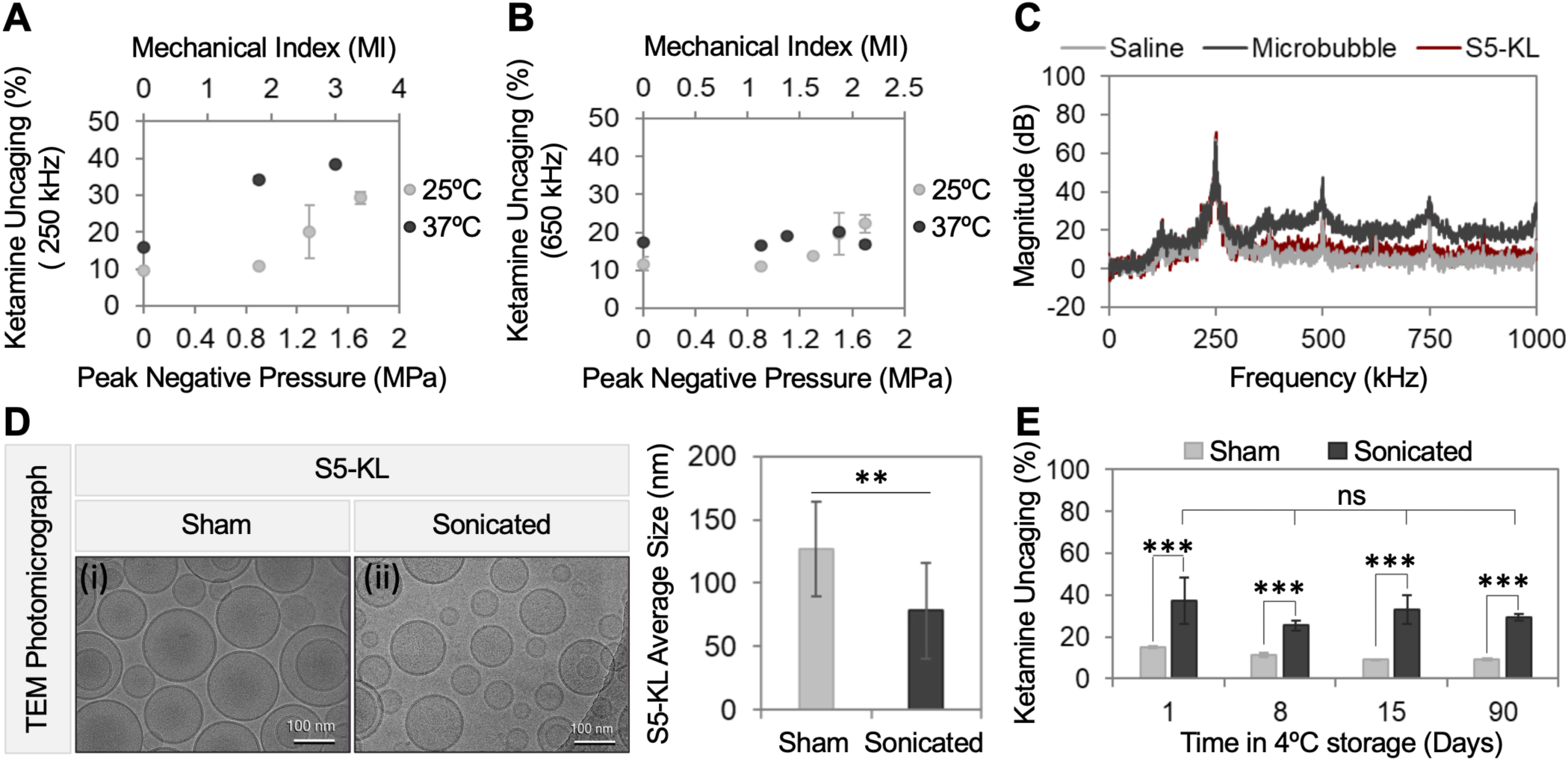
Investigating the mechanism of ultrasonic drug uncaging from AALs. Ketamine uncaging from S5-KL with ultrasound application (25% duty cycle, PRF 5 Hz, 60 s) with varying peak negative pressure and MI, at 25 °C or 37 °C with (**A**) 250 kHz and (**B**) 650 kHz center frequency. Increased ketamine release observed at 250 kHz across different pressures/MIs and temperatures compared to 650 kHz. (**C**) Magnitude of the received echo spectrum following ultrasound applied to saline (light gray), gas-filled microbubbles (black) or S5-KL (red line) as a function of frequency following sonication with a center frequency of 250 kHz, 0.9 MPa peak negative pressure, and 25% duty cycle indicates no subharmonic, ultraharmonic, or broad band acoustic scatter indicative of cavitation with S5-KL, with no difference in sonication of saline versus S5-KL. In contrast, sonication of microbubbles yields broad band, subharmonic, and ultraharmonic signal induction, indicating cavitation. (**D**) Representative Cryo-EM images show spherical shape of the S5-KL liposomes and that the size distribution of the particles decreased with sonication. (**E**) Stability of S5-KL for both drug retention while exposed to plasma and for ultrasound-induced release across 90 days in 4 °C storage. Data presented as mean ± SD of three or more independent experiments. Comparisons between two groups were performed by two-tailed Student’s t-tests; ns= non-significant, *p < 0.05, **p < 0.01 and ***p < 0.001.

### AALs as a generalized platform for ultrasonic drug uncaging

To confirm that these results were not somehow specific to the particular drug used in these initial experiments, we prepared liposomes with or without 5% additional sucrose internally, and loaded with several other agents: lidocaine, bupivacaine, and ropivacaine. These liposomes show similar size, polydispersity, and zeta potential as the ketamine-loaded liposomes (Table 1). They showed differential drug loading rates that likely reflect the specific chemical characteristics of each drug, such as their logP and pKa, as they relate to the ammonium sulfate active loading mechanism (Table 1). Each of these other liposomes showed increased ultrasonic drug uncaging with the formulations incorporating added sucrose internally, while formulations without added sucrose showed no significant or minimal drug uncaging with ultrasound (Fig. 3).

**Fig. 3.**
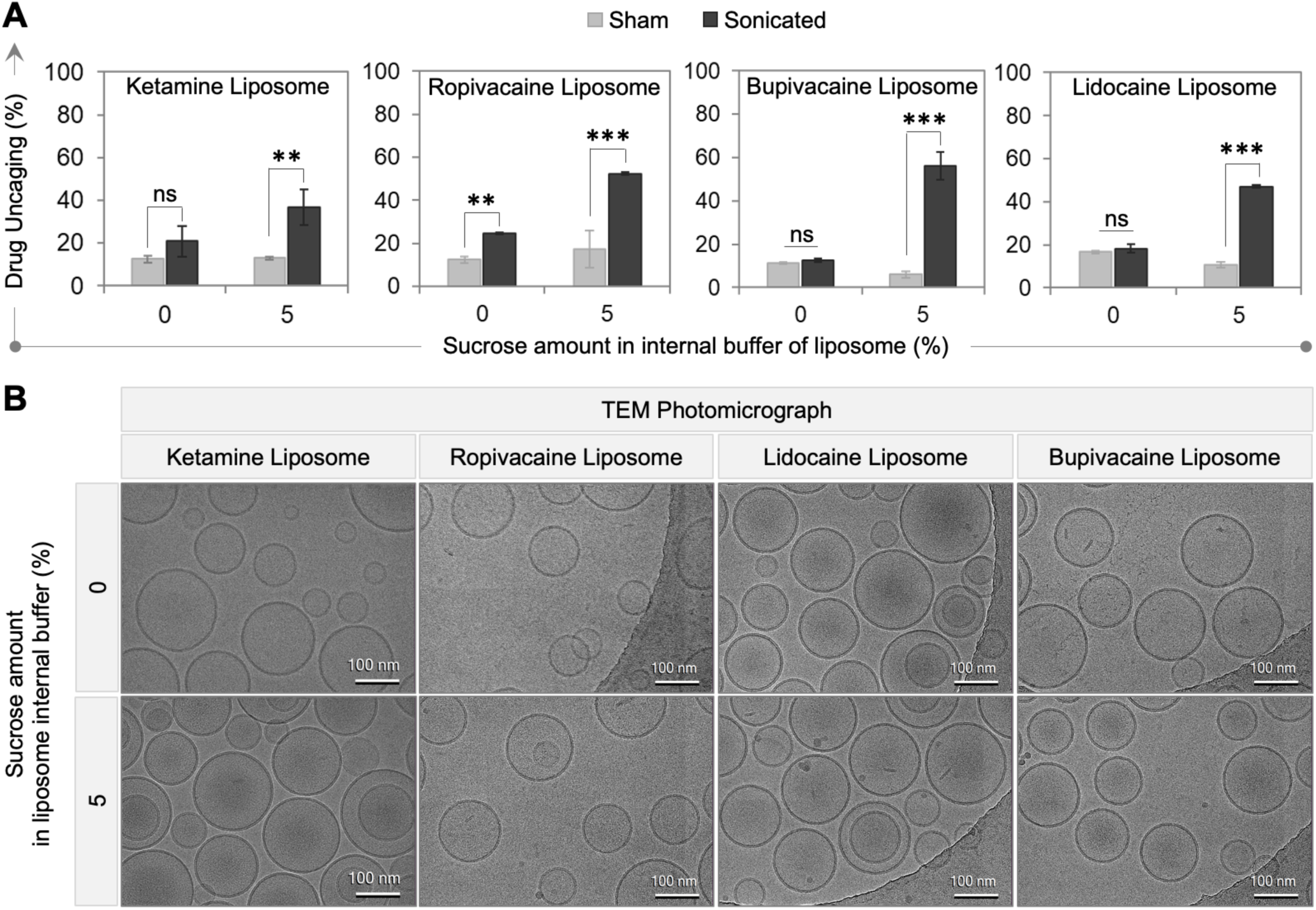
AALs are a platform for ultrasonic uncaging of a variety of drugs. (**A**) To confirm the generalizability of the concept, several agents – ketamine, ropivacaine, bupivacaine and lidocaine – were loaded into the liposome with or without 5% additional sucrose internally. Each drug loaded formulation shows higher in vitro ultrasonic drug uncaging with sucrose incorporated versus the no sucrose control formulations. Data presented as mean ± SD of three independent experiments. Comparisons between two groups were performed by two-tailed Student’s t-test; ns= non-significant, *p < 0.05, **p < 0.01 and ***p < 0.001. (**B**) Cryo-EM images of each formulation confirm their spherical morphology, without a specific morphologic change with sucrose incorporation. Ketamine data partly repeated from Figs. 1 and 2.

### In vivo pharmacodistribution and ultrasonic drug uncaging of AALs

With these in vitro characterizations demonstrating in vitro stability and ultrasonic drug uncaging efficacy, we next characterized their performance in vivo. Following a bolus intravenous injection, the blood drug concentration showed slower blood clearance pharmacokinetics with an acoustomechanically activatable liposome (AAL) versus a dose-matched administration of a free unencapsulated drug formulation (Fig. 4A). Notably, there were also higher drug metabolite blood levels with the AAL formulation compared to the unencapsulated drug formulation. This behavior was reflected in the biodistribution at 1 hr post bolus administration, in which the solid organs showed increased accumulation of the drug with the free unencapsulated form administration compared to the AAL formulation (Figs. 4B, S1). The combination of the higher blood drug metabolite concentration yet lower unmetabolized drug accumulation in the solid organs with the AAL formulation suggests that some of the blood clearance of the drug with the AAL formulation is likely due to the sequestration and metabolization of the whole particle by end organs like the liver. Further, it is noted that this end organ sequestration and metabolism could contribute to some degree of nonspecific leak of the drug into circulation, and may be less present a mechanism in humans vs. rodents given the lower hepatic blood flow and hepatic mass (normalized by body weight) in humans.

**Fig. 4.**
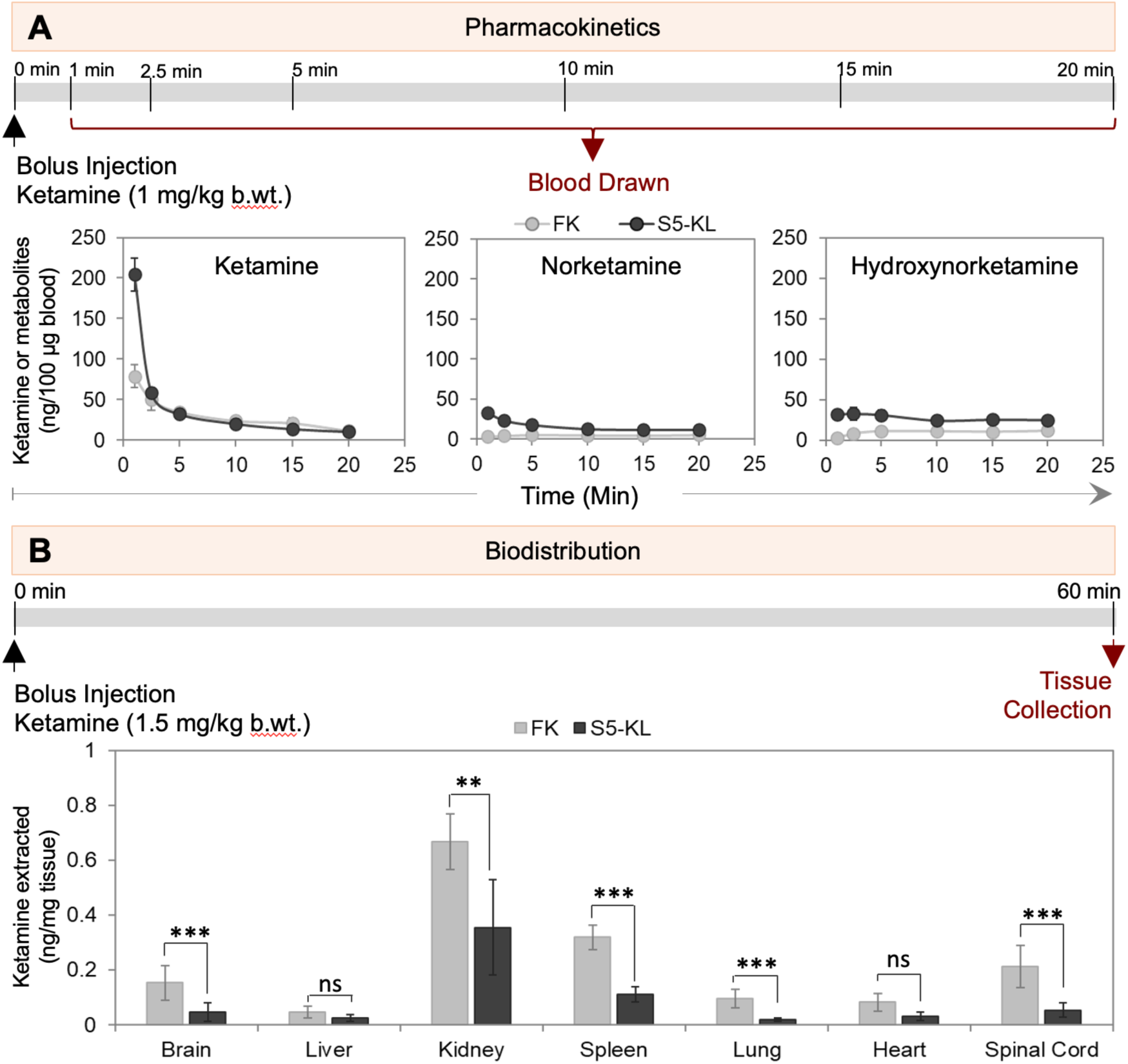
In vivo AAL pharmacokinetics and biodistribution. (**A**) Ketamine quantification in blood samples collected at different timepoints after 1 mg/kg body weight intravenous bolus injection of free, unencapsulated ketamine-HCl (FK) or S5-KL (n=4 adult rats each) shows slower blood clearance with S5-KL compared to FK for ketamine itself and its major metabolites norketamine and hydroxynorketamine. (**B**) Ketamine quantification in different organs after 60 min following 1.5 mg/kg body weight intravenous bolus injection of FK or S5-KL shows higher solid organ drug accumulation with ketamine-HCl (FK) compared to the ketamine-loaded AAL (S5-KL; n=4 adult rats each). Data presented as mean ± SD. Comparisons between multiple groups performed by one way analysis of variance (ANOVA), ns= non-significant, *p < 0.05, **p < 0.01 and ***p < 0.001.

To assess the differences induced of in vivo pharmacodistribution with ultrasonic drug uncaging with AALs, in adult rats, we used solid-phase microextraction (SPME) to measure the amounts of ketamine delivered to the brain either with a free unencapsulated drug infusion or an AAL infusion, with or without ultrasound application to the brain. Ultrasound (or sham administration) was applied to a frontal cortical and hippocampal region unilaterally during the latter 2.5 min of a 5 min intravenous infusion of either free, unencapsulated ketamine-HCl or ketamine-loaded liposomes (Fig. 5A). After the stop of infusion and sonication, SPME was used to sample both the sonicated brain and a contralateral control region. A venous blood sample was taken at the end of the SPME sampling period. The blood samples showed higher blood levels of ketamine with each liposomal formulation compared to the free drug infusion, confirming our blood pharmacokinetics results following bolus administration (Fig. 5B, 4A). Nonetheless, despite the higher blood concentrations of ketamine with liposomal formulation administration, the ketamine brain levels seen either with sham ultrasound or in the contralateral control region were less than half of that seen following a free unencapsulated ketamine infusion (Fig. 5C). While ultrasound application did not measurably alter the pharmacodistribution of ketamine following a free ketamine infusion, ultrasound application yielded significantly higher brain ketamine levels with the liposomal infusions compared to both the sham ultrasound condition and to the contralateral non-sonicated brain that was separated by 5 mm (Fig. 5A,C). Notably, this up to 5 mm of spatial localization of the uncaging effect seen in this characterization is just beyond the full-width at half-maximum of the 250 kHz ultrasound beam used in this application, confirming that this uncaging is limited spatially by the applied ultrasound field. Furthermore, significantly higher brain ketamine levels were seen with sonication with liposomes made with additional sucrose internally compared to liposomes without added sucrose, confirming that this relatively simple formulation manipulation increases the ultrasound sensitivity of the system both in vitro and in vivo. Notably, despite higher ultrasonic drug uncaging efficacy in vitro, no significant difference was seen in vivo between liposomes made with 10% added sucrose internally, compared to those made with 5% added sucrose, likely reflecting the relative instability in plasma of the 10% added sucrose formulation seen in vitro (Fig. 1E). Importantly, histological assessment for morphological tissue damage, neuronal degeneration, microglial activation, and astrocytic activation and gliosis yielded no evidence of brain parenchymal injury seen with AAL-mediated ultrasonic drug uncaging (Figs. 6, S2).

**Fig. 5.**
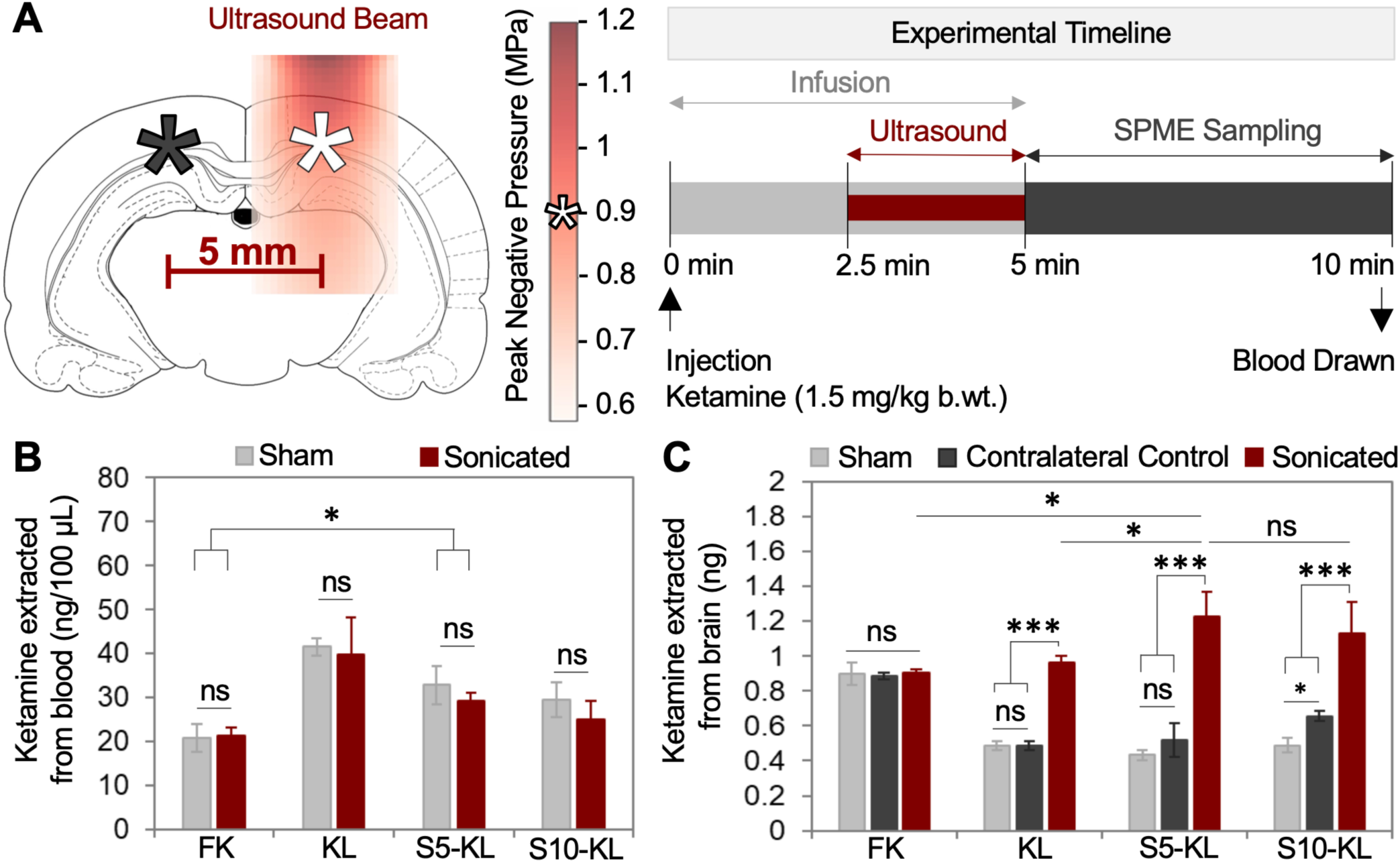
In vivo uncaging of ketamine-loaded AALs. (**A**) Schematic and timeline for the in vivo pharmacodistribution study. Ultrasound (250 kHz, 0.9 MPa est. peak in situ neg. pressure at the sampling site center (white asterisk), 25% duty cycle, 5 Hz PRF; or sham) was applied to a frontal cortical/hippocampal brain region unilaterally during the latter 2.5 min of a 5 min intravenous infusion of 1.5 mg/kg body weight of either free unencapsulated ketamine-HCl (FK), or ketamine liposomes without (KL) or with 5% (S5-KL) or 10% (S10-KL) added sucrose in the internal buffer. Sonication beam map constructed based on hydrophone beam mapping of the transducer with intensity scaled according to the applied intensity derated for rat skull insertion loss. After sonication, solid phase microextraction (SPME) was used to sample both the sonicated brain (white asterisk) and a contralateral control region (black asterisk) separated 5 mm apart. A venous blood sample was taken at the end of the SPME sampling period. Group N: FK Sham (8), FK Sonicated (2), KL Sham (3), KL Sonicated (4), S5-KL Sham (5), S5-KL Sonicated (6), S10-KL Sham (3), S10-KL Sonicated (4). (**B**) Ketamine quantification in blood shows higher levels of drug with the liposomal formulations (KL, S5-KL, & S10-KL) compared to the free drug (FK) infusion. (**C**) Ketamine quantification shows no alteration of the concentration of drug in brain with ultrasound application following free drug infusion. With each liposomal formulation infusion, ultrasound application significantly increased the drug concentration in the sonicated brain region compared to the contralateral or sham controls, with increased uncaging seen with sucrose incorporation. Data presented as mean ± SD. Comparisons between two groups was performed by two-tailed Student’s t-test, and that among multiple groups by one way analysis of variance (ANOVA); ns= non-significant, *p < 0.05, **p < 0.01 and ***p < 0.001.

**Fig. 6.**
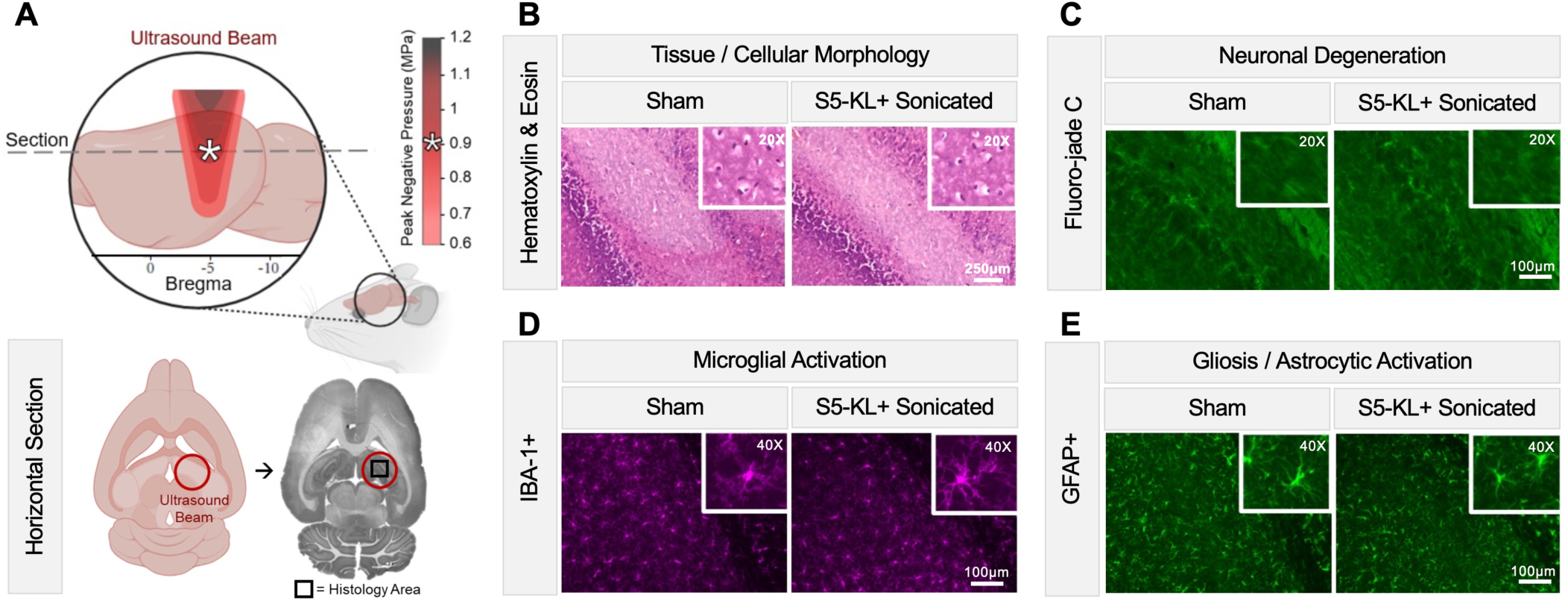
Ultrasonic drug uncaging in the brain with AALs is safe. Histological and immunohistochemical analyses of brain tissue 72 hrs after sonication with AALs (N=3/group) show no indication of parenchymal damage in comparison to healthy controls. (**A**) Schematic representation of the rat brain depicting the region of ultrasound sonication and histological sampling. (**B**) Representative magnified images (4x, large, and 20x, inset, zoom) of hematoxylin and eosin staining shows no alteration in tissue and cellular morphology. (**C**) Representative magnified images (10x, large, and 20x, inset, zoom) of Fluoro-Jade C staining for neurodegeneration shows no change in neuronal degeneration (FJ-C positive cells, green). (**D**) Representative magnified images (10x, large, and 40x, inset, zoom) of IBA-1 immunostaining indicate no change in microglial activation (IBA-1 positive cells, purple). (**E**) Representative magnified images (10x, large, and 40x, inset, zoom) of GFAP immunostaining shows no indication of gliosis or change in astrocytic activation (GFAP positive cells, green).

### Functional efficacy of AAL-mediated ultrasonic drug uncaging

Finally, to assess the functional efficacy and generalizability of AAL-mediated ultrasonic drug uncaging, we assessed whether uncaging with an AAL loaded with a different drug could induce a behavioral effect. Specifically we tested whether uncaging an AAL loaded with a local anesthetic could be used to induce anesthesia of a peripheral nerve, as a form of a noninvasive needle-less nerve block. Adult rats were administered a 5 min intravenous infusion of either saline or a ropivacaine-loaded AAL (S5-RL). With 2.5 min of pulsed sonication during the latter half of the infusion applied to the sciatic nerve of one limb, a dense anesthesia to a mechanical (von Frey fiber) stimulus was observed only in the sonicated limb of the ropivacaine-AAL administered animals, with this anesthesia lasting at least 4 hrs following infusion and sonication (Fig. 7C). No such change was seen in either limb of the saline infused animals or in the contralateral non-sonicated limb of the ropivacaine-loaded AAL treated animals (Fig. 7B,C). Notably, in rats receiving this same dose of ropivacaine-loaded AALs, no significant electrocardiographic abnormalities or visible intolerances were noted, confirming the safety of this approach (Fig. 7D-F).

**Fig. 7.**
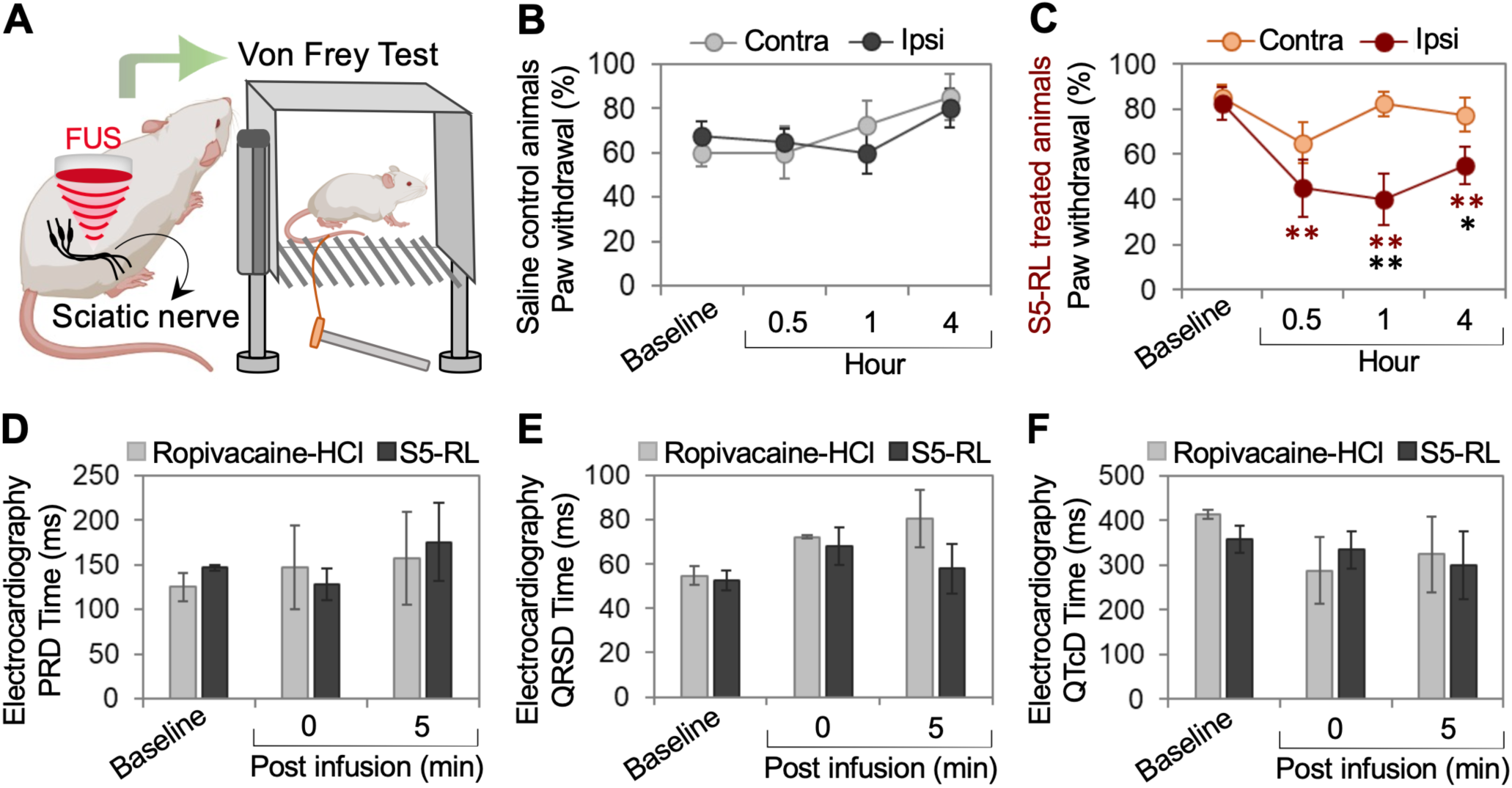
Functional efficacy of in vivo AAL-mediated ultrasonic ropivacaine uncaging. (**A**) Adults rats (n=8/group) received a 5 min intravenous infusion of saline or a ropivacaine-loaded AAL (S5-RL, 5 mg/kg), with focused ultrasound (FUS; 250 kHz, 25% duty cycle, 5 Hz PRF, 0.9 MPa est. in situ peak neg. pressure) applied to the sciatic nerve of one limb for 2.5 min during the latter half of the 5 min infusion. Frequency of paw withdrawal to a mechanical stimulus (von Frey test) was assayed before and after infusion and sonication. (**B**) In saline and FUS treated animals, no significant difference in von Frey test results were seen across time or between the limb ipsilateral (ipsi) to sonication vs. the limb contralateral (contra) to sonication. (**C**) In SR-5L and FUS treated animals, significant differences on von Frey testing were seen in the limb ipsilateral (ipsi) to sonication at time points after sonication and infusion vs. the baseline (red asterisks), as well as at certain time points between the limb ipsilateral (ipsi) to sonication vs. contralateral (contra) to sonication (black asterisks). (**D-F**) No significant differences were seen of key electrocardiographic parameters with a 5 min intravenous infusion of 5 mg/kg ropivacaine-HCl or S5-RL (n=4/group). Comparisons between multiple groups performed by two way analysis of variance (ANOVA); ns= non-significant, *p < 0.05, **p < 0.01 and ***p < 0.001.

## DISCUSSION

With these results, we have demonstrated a system for ultrasonic drug uncaging that can be loaded with varied drugs with high drug loading, minimal drug release without ultrasound, and effective drug release with pulsed low-intensity ultrasound both in vitro and, importantly, in vivo for multiple drugs of interest. The ultrasound-induced drug release mechanism with this system does not necessitate medium heating or cavitation, and shows a frequency response characteristic indicating a mechanical type interaction, with significant ultrasound-induced drug release achievable with pulsed ultrasound with low intensities that are within FDA and AIUM guidelines for safe ultrasound application (*46, 47*). Notably in vivo, the uncaging effect could be localized to within 5 mm of the center of the applied ultrasound field, just beyond the full-width at half-maximum of the 250 kHz ultrasound field used in these experiments. Additionally, we found that AAL-mediated ultrasonic drug uncaging could have behavioral functional relevance, as demonstrated with ultrasonic ropivacaine uncaging yielding a long-lasting noninvasive needle-less peripheral nerve block. Crucially, we have been able to achieve these features with formulation constituents that are commonly used in clinical formulations as validated pharmaceutical excipients (*48, 49*), with the simple disaccharide sucrose being the main additional ingredient yielding increased ultrasound sensitivity in this current implementation of acoustomechanically activatable liposomes (AALs).

By our design, it seems that additional sucrose in this formulation increases the acoustic impedance of the inner core droplet of the liposome relative to the surrounding medium, and that this acoustic impedance difference permits an ultrasound interaction that increases drug leak across the liposome membrane. We further postulate that the osmotic pressure difference across the lipid membrane further accelerates drug release following the initial acoustic interaction induced lipid membrane leakiness (*46, 50, 51*). While we have focused our attention on implementing this system using common pharmaceutical excipients to maximize clinical translatability, an implication of this acoustomechanical release mechanism is that an even further ultrasound-responsive system could be achieved by replacing sucrose with equiosmolar amounts of a different substance that even further shifts the acoustic impedance away from that of plasma.

In this effort, we sought to also maintain clinical translatability of this system. Indeed, the manufacturing methods for liposome production used here are compatible with Good Manufacturing Practices (GMP) and the formulation ingredients are each validated excipients in the FDA Inactive Ingredients Database (*39*), together indicating a ready path to clinical translation of this system. Additionally, this formulation is stable for months in refrigerated storage, a decided practical advantage over prior formulations that necessitated frozen storage or fresh preparation (Table 2). In addition, the ultrasound intensities sufficient for drug release with this system have a mechanical index within the FDA and AIUM guidelines of safe ultrasound application (*45*); indeed no evidence of parenchymal damage nor of visible or electrocardiographic intolerance with AAL uncaging was observed in our in vivo experiments. Further, this ultrasound protocol is readily achievable using current clinically utilized therapeutic focused ultrasound systems (*52, 53*). Finally, these AALs enable targeted drug delivery to within a few millimeters resolution in the rat brain following an intravenous infusion, a notable practical advantage over systems necessitating direct intraparenchymal injection of the drug-loaded formulation.

**Table 2.** AAL performance and practicality compared to benchmark ultrasonic drug uncaging systems. N/A: Not available.

| Ultrasonic drug uncaging systems | In Vivo Use and Performance |  |  | Clinical Translatability and Practicality |  |  | Notes | Ref. |
| --- | --- | --- | --- | --- | --- | --- | --- | --- |
|  | Mechanism | Ultrasound parameters | Spatial localization; Drug delivery effect size | Excipients not in FDA IID or active clinical use | Drug load | Storage, Stability |  |  |
| Acousto-mechanically activatable liposomes | Mechanical, non-cavitation | <u>Frequency:</u> 250 kHz<br><u>Duty Cycle</u> 25%<br><u>Peak Pressure:</u> 1.8 | <5 mm @ 250 kHz<br>3.1x increase of ketamine in brain | None | 3 mg/mL ketamine | 4 °C storage; stability >3 mo |  |  |
| Polymeric liquid perfluorocarbon nanodroplets | Vaporization/ Cavitation | <u>Frequency:</u> 650 kHz-3 MHz<br><u>Duty Cycle</u> 1-20%<br><u>MI:</u> 1.5-2.25 | <2 mm @ 650 kHz (functional assay; drug delivery N/A)<br>Drug delivery effect: N/A | Polymer shells, perfluorocarbon cores | 1 mg/mL propofol in (12, 21) | -80 °C storage; stability >1mo for (12, 21) | No cavitation seen in (12, 21) | (12, 21, 65–68) |
| Lipid liquid perfluorocarbon nanodroplets | Vaporization/ Cavitation | <u>Frequency:</u> 580 kHz, 1.66 MHz<br><u>Duty cycle:</u> 1%<br><u>Pressure:</u> 0.8 MPa (580 kHz) or 2.3-2.8 MPa (1.66 MHz) | <1 mm for a dye (drug delivery N/A) @ 1.66 MHz<br>Drug delivery effect: N/A | Perfluorobutane | 25 µg/mL pentobarbital in (8) | Prepared fresh; Storage and stability N/A |  | (8, 13, 20, 69) |
| Temperature-sensitive liposomes | Heating | <u>Frequency:</u> 1 MHz<br><u>Duty Cycle</u> 100%<br><u>Intensity:</u> N/A (heating to >40 °C) | Spatial localization in vivo: N/A<br>3.7x increase of doxorubicin in liver in (71) | None | 3.4 mg/mL doxorubicin in (70) | 4 °C storage; stability >7 days |  | (70, 71) |
| Sonodynamic liposomes | Sonodynamic oxidation induced membrane leak | <u>Frequency:</u> 1 MHz<br><u>Duration:</u> 10 min in (34)<br><u>Duty Cycle:</u> 100%<br><u>Peak Power Density:</u> 3 W/cm<sup>2</sup> in (34); 0.2-0.3 W/cm<sup>2</sup> in (72) | Spatial localization in vivo: N/A<br>Drug delivery effect: N/A | Protoporphyrin IX; Porphyrin | API conc. N/A; 63-92% loading efficiency | 4 °C storage; stability >1 month in ref. 14 | Direct parenchymal agent injection needed in (34) | (34, 72) |
| Microbubble-Liposome Conjugates | Induced aggregation then cavitation | <u>Frequency:</u> 2.5 MHz<br><u>Peak pressure x time:</u> 0.075 MPa x 1000 ms then 0.188 MPa x 1000 cycles x 100 Hz PRF for 90 ms then 300 ms off | Spatial localization in vivo: N/A<br>Drug delivery effect: N/A | DSPE-PEG5000-Maleimide; DSPE-PEG5000-Thiol; Perfluorobutane | 200 ng/5-6 mL muscimol for in vivo dose | Prepared fresh; Storage and stability N/A |  | (9) |
| Carbon dioxide loaded liposomes | Cavitation | <u>Frequency:</u> 237 kHz or 1.57 MHz<br><u>Pressure:</u> 0.67 MPa or 0.24 MPa | Spatial localization in vivo: N/A<br>Drug delivery effect: N/A | None | API conc. N/A; 5-46% loading efficiency | 4 °C storage; Stability N/A | No in vivo use | (73) |

While an ammonium sulfate based active loading scheme was demonstrated here, the strategy of incorporating elements to increase the acoustic impedance of the liposome core to increase its ultrasound responsiveness should be applicable to any liposome with any internal core buffer and loading battery (*48, 54, 55*). Future efforts will explicitly demonstrate the utility of these acoustomechanically activatable liposomes for ultrasonic uncaging of other pharmaceutical agents that may be so differently encapsulated. Of note, there was a nonzero rate of drug leak without ultrasound both in vitro and in vivo. While the in vivo ultrasound-induced uncaging effect size seen with these AALs would be sufficient to identify a relevant therapeutic index for each drug tested, and are in line with or exceed other benchmark systems that have been so characterized (Table 2), this baseline nonspecific delivery potentially reflects the lack of internal precipitation seen with these particular drug-loaded liposomes, and potentially the metabolism of the AAL contributing to drug leak during the metabolism. Notably, this latter effect may be less contributory in humans given the relatively lower rate of hepatic blood flow and metabolism in humans compared to rats. Future efforts will attempt to reduce the baseline release rate of AALs while further maximizing ultrasound-responsiveness by including materials that increase internal drug binding while further shifting the liposome core acoustic impedance.

## MATERIALS AND METHODS

### Materials

All the chemicals and reagents used in the study were of the highest purity grade with sourcing as follows. Lipoid: Hydrogenated Soy Phosphatidylcholine (HSPC) (Cat no: Lipoid S PC-3) and [N-(methoxypolyethyleneglycol-2000)-1,2-distearoyl-sn-glycero-3-phosphoethanolamine, sodium salt] (DSPE-PEG 2000) (Cat no: PE18:0/18:0 – PEG 2000). Evonik: Cholesterol (PhytoChol Inject HU). Stanford University Environmental Health & Safety: Ketamine-HCl injectable solution (100 mg/mL). Astatech: Ropivacaine-HCl (Cat no. 42084) and Bupivacaine-HCl (Cat no. 42055). Hiemdia: Ammonium Sulfate (Cat no: PCT0003). Fisher Scientific: Absolute Ethanol (200 proof) (Cat no: BP2818-100), Sucrose (Cat no: S5-500), HPLC and LC/MS grade Water, Methanol, Acetonitrile, 2-Propanol, Formic Acid. Sigma Aldrich: Lidocaine-HCl (Cat no. L5647), HEPES buffer solution (Cat no: 83264-500ML-F) and L-Histidine monohydrochloride monohydrate (Cat no: H8125). Repligen: TFF filters (C02-S05U-05-N (SN: 20020493-03/21-057). Lampire Biological Laboratories: Male Canine Plasma (Cat no. 7302009). Cytiva: PD-10 column Sephadex G-25 M (Cat no. 17085101), Sephacryl S-500 (Cat no. 1706130). Milli-Q water was used to prepare all the buffers. Millipore Sigma: Cerilliant® certified standard solutions of ketamine hydrochloride, ketamine-D4 hydrochloride, norketamine hydrochloride, norketamine-D4 hydrochloride, and hydroxynorketamine hydrochloride; Supel™ BioSPME 96-Pin Devices (product number 59680-U).

### Liposome production

An ammonium sulfate remote loading scheme was utilized for each reported liposome, following published protocols (*56*). Briefly, small unilamellar vesicles were prepared by dissolving lipid components (HSPC:DSPE-PEG2000:cholesterol 52.8:42.3:4.8 molar ratio) in heated ethanol and then diluting the mixture to 10% ethanol with 250 mM ammonium sulfate alone or with 5-10% (by weight) sucrose, 5% glucose, or 73 mM NaCl depending on the experiment. Then the solution was extruded (Avestin LF-50 with 200 nm pore polycarbonate Whatman filter at 65-70 °C) ten times to generate the unilamellar liposomes. Then, the sample was processed with tangential flow filtration (5x 5-fold dilution/reconcentration) against a buffer of 10 mM HEPES, 145 mM NaCl (pH 7.4) with 0, 5, or 10% sucrose depending on the internal buffer osmolarity, to generate a transmembrane ammonium gradient. For remote loading, the liposomes were diluted 10-fold and then the drug was added as a hydrochloride salt to a 1 mg/ml concentration and the mixture was heated to 55 °C for 1.5 h. Then TFF (4x 5-fold dilution/reconcentration) was repeated against a buffer of 10 mM histidine (pH 7.4) with 10 or 15% sucrose. Finally, the samples were filtered through a sterile 220 nm PVDF syringe filter inside a sterile hood and the sterilized filtrate were dispensed into aliquots, sealed, and stored at 4 °C for future use.

### Sizing & Zeta potential

The Z-average diameter, polydispersity index (PDI) and zeta potential of liposomes were measured with a Malvern Zetasizer Nano ZS90 (Malvern, United Kingdom), reading a 1:100 diluted sample of liposomes. All measurements were carried out in triplicates with the auto mode with the following settings: 20 to 30 runs per measurement, temperature = 25 °C, refractive index of Lipid = 1.5.

### Cryo-electron microscopy (cryo-EM)

Cryo-EM was performed on a ThermoFisher Scientific Glacios™ Transmission Electron Microscope (Cryo-TEM) operating at 200 kV. 3 μL of purified samples were applied on glow-discharged (40 sec, 15 mA, PELCO easiGlow™, TED PELLA Inc.) holey carbon grids (Quantifoil Cu R1.2/1.3, 200 mesh). The grids were blotted using a Vitrobot Mark IV (FEI) with 3 sec blotting time, blot force 2 at 22°C in 100% humidity, and plunge-frozen in liquid ethane. The plunge frozen grids were clipped and loaded into the Glacios EM. The specimen was examined at 36,000x magnification and images were recorded with a pixel size of 1.15 Å using SerialEM software (University of Colorado, Boulder) and a K3 direct electron camera (Gatan). Autofocusing performed with defocus range set to 1.5-3 microns.

### High-Performance Liquid Chromatography (HPLC)

HPLC was carried out using an Agilent 1260 Infinity II HPLC System (Agilent Technologies, San Jose, CA) equipped with a multiple wavelength UV detector (G7165A) using Agilent OpenLab CDS software. The chromatographic separation was executed at ambient room temperature using a reverse-phase Agilent 5 HC-C18(2) column (250 mm × 4.6 mm, 5 µm particle size) (Cat no. 588905-902) equipped with an Agilent 5 HC-C18(2) guard column (12.5 mm × 4.6 mm, 5 µm) (Cat no. 520518-904). Water and acetonitrile with 0.1% trifluoroacetic acid were used as the mobile phase. A linear gradient elution was used with detection wavelength of 270 nm for ketamine, and 263 nm for lidocaine, bupivacaine, and ropivacaine.

### Drug concentration quantification

Samples were diluted 1:15 in HPLC grade Methanol and measured using HPLC according to the USP method for each drug.

### Unencapsulated drug quantification

First, 2.5 mL of 1:4 diluted liposomes were loaded onto a PD10 size exclusion column. Then, buffer was added. The initial 4.5 mL eluted fraction contained the liposomes and the next 9 mL of elute contained the unencapsulated drug. The drug concentration in each fraction was determined by HPLC and the unencapsulated fraction determined as the percent in the 2^nd^ eluted fraction versus the total in the 1^st^ and 2^nd^ fractions.

### In vitro ultrasonic drug uncaging

First 200 μl of liposomes diluted 1:4 in canine plasma were loaded into 0.2 mL thin walled PCR tubes placed in a custom 3D printed holder held at the focus of a 250 or 650 kHz hydrophone-calibrated focused ultrasound transducer. Degassed water of either 25 or 37 °C was used to couple the transducer to the sample tube. Then ultrasound (60 s, 25% duty cycle, 5 Hz PRF, varying peak pressure) was applied. For analysis, each applicated was repeated 5x and these fractions were collected together into 1 mL. Liposomes were separated from unencapsulated drug by a homemade Sephacryl S-500 column, with PBS as the elution buffer, collecting the first 5.5 mL of elute as the liposome fraction and the next 8 mL as the uncencapsulated drug fraction. Drug concentration in each elute was quantified by HPLC.

### Speed of sound measurement

A clean 20-gallon fish tank was filled with deionized water and degassed overnight. A 650 kHz 30 mm aperture f1.0 focused ultrasound transducer (Sonic Concepts Inc, Bothell, WA), a 35.6 cm long PVC cylinder with a 1.5-inch diameter, and a capsule hydrophone (Onda Corp, Sunnyvale, CA) were placed in a row underwater. Both devices were linked to an oscilloscope (Keysight Technologies, Santa Rosa, CA) to view the time of flight between the transducer and the hydrophone. The PVC pipe was wrapped in an ultrasound-compatible plastic probe cover and sealed with O-rings to create a separate internal fluid compartment. First, the pipe was loaded with 37 °C DI water, and the ultrasound pulse arrival time was used as a reference, along with the known speed of sound in DI water at 37 °C, for subsequent measurements. Each buffer was sequentially loaded after heating, and the difference in pulse arrival time was recorded. A temperature measurement after each run confirmed minimal heat loss. The differences in pulse arrival time due to different speeds within the length of the pipe were translated into speeds of sound of the various buffers. Finally, measurements of density at 37 °C gave the acoustic impedance value as Z = (speed of sound)*(density) for each buffer.

### Cavitation assessment

The same setup used for in vitro ultrasonic drug uncaging was applied with a capsule hydrophone (ONDA, Sunnyvale, CA) placed beside the PCR tube perpendicular to the ultrasound focus Z-axis. The emitted and scattered signals from the samples inside the PCR tube was detected by the hydrophone, recorded, and processed by Fourier transform by the oscilloscope, using the first 2 ms of signal.

### Animal preparation

All animal experiments were carried out in accordance with the Stanford IACUC and Administrative Panel on Laboratory Animal Care (APLAC). Male Long-Evans rats 7-10 weeks old, body weight 250-400 g (Charles River Laboratories, Wilmington, MA; or Envigo, Indianapolis, IN) were used in all in vivo studies. Rats were anesthetized under 2% isoflurane for the entire experiment and fixed into a stereotaxic frame. They were administered 2 mL of saline subcutaneously and kept on a heating pad at 37 °C. Ketamine-loaded liposomes, ketamine-HCl, or saline vehicle were administered by tail vein catheter. A 250 kHz focused ultrasound transducer powered by an amplifier (240L, E&I Ltd, Rochester, NY) was utilized in all in vivo ultrasound experiments. Skull attenuation was accounted for and calculated based on weight to achieve the desired in situ pressure (*57*).

### In vivo ultrasonic drug uncaging

Before animal experiments, BioSPME pins were carefully detached from their housing, cleaned with 2:1:1 v/v/v methanol:acetonitrile:isopropanol, then preconditioned overnight in 1:1 v/v methanol:water, then transferred to water until sampling. After anesthesia induction, the rat dorsal scalp was exposed and 2- mm burr holes were drilled bilateral into the skull for SPME pin insertion (5 mm posterior to bregma, +/- 2.5 mm lateral to midline). A durotomy was performed with a 32g needle. The ultrasound transducer was positioned directly above the right burr hole via a 3-axes positioning system (ThorLabs, Newton, NJ), and ultrasound gel was used for coupling. Drugs were infused over 5 minutes intravenously via a tail vein catheter with an infusion pump (World Precision Instruments, Sarasota, FL). Focused ultrasound was applied after 2.5 minutes of drug infusion for 2.5 minutes (250 kHz, 0.9 MPa estimated peak in situ pressure, 5 Hz PRF, 25% duty cycle). For sampling, the SPME pins were loaded into a custom-designed, 3D-printed stereotaxic holder and the SPME pins were inserted 3 mm ventrally into the brain via the burr holes (SPME absorptive medium centered at ∼2.5 mm ventral to bregma). Following 5 min exposure to the rat brain tissue to allow drug/metabolite diffusion (*58–62*) to the device, the SPME pins were retrieved carefully, washed for 5 seconds via static immersion in water, and desorbed into 50 µL of MeOH:H_2_O (9:1 v/v) containing 1% formic acid and deuterated internal standards. Following 30 min of desorption with agitation at 1500 rpm at room temperature, the desorbed aliquots were quantified by LC-MS/MS analysis employing an instrumental calibration curve.

### Pharmacokinetics & Biodistribution

For the pharmacokinetics experiment, 8 adult rats (N=4 each group), following anesthesia, received an intravenous bolus dose of 1 mg/kg of ketamine as either free ketamine-HCl or ketamine liposomes. Then, blood samples (approximately ∼400 μL) were collected by tail snipping into Eppendorf tubes pre-loaded with 50 μL Na_2_EDTA at 1, 2.5, 5, 10, and 15 min. For the biodistribution experiment, another 8 adult rats (N=4 each group) received either free or liposomal ketamine as an intravenous bolus dose of 1.5 mg/kg. Rats were sacrificed at 1 hour from the time of administration and perfused with 1x phosphate buffer (PBS) via transcardial perfusion to remove blood from the systemic circulation. Afterward, the tissues, including the brain, liver, kidney, spleen, lung, heart, and spinal cord, were collected and stored at -80 °C.

The solid organ samples were homogenized in equal-weight volume of 1x phosphate buffer (PBS) using a handheld homogenizer. 100 μL of tissue homogenate was aliquoted from each organ in triplicate per organ. Similarly, 100 μL of blood sample from each time point was aliquoted into triplicates. Deuterated internal standards (ketamine-D4 and norketamine-D4) were added to each blood and tissue homogenate sample. Protein precipitation was carried out with 500 μL of cold acetonitrile. The samples were subjected to an hour of extraction using a mechanical shaker, followed by centrifugation at 12000 rpm and 4°C for 10 minutes. The resulting supernatant was collected and after evaporation of the organic solvent, the sample was reconstituted with 100 μL of MeOH:H_2_O (9:1) containing 1% formic acid and quantified by LC-MS/MS analysis using an instrumental calibration curve.

### LC-MS/MS analysis

The LC-MS/MS analysis was conducted using an Agilent 1290 Infinity LC System coupled to a 6490 Triple Quadrupole LC/MS System with iFunnel technology (Agilent Technologies, San Jose, CA). The MS/MS analysis was executed in the positive mode, utilizing Agilent Jet Stream technology for electrospray ionization (AJS-ESI), operating under the conditions of selected reaction monitoring (SRM). The fragmentor voltage of 380 V and a cell accelerator voltage of 5 V were consistently applied for all SRM transitions. Data acquisition and processing were performed using Agilent MassHunter Workstation Data Acquisition and MassHunter Workstation Quantitative Analysis software (Agilent Technologies, San Jose, CA), respectively. The chromatographic separation was using Agilent ZORBAX RRHD Eclipse Plus C18 column (5 cm × 2.1mm, 1.8 µm particle size; cat no. 959757-902) equipped with an Agilent ZORBAX RRHD Eclipse Plus C18 guard column (2.1 mm, 1.8 µm; cat no. 821725-901). Water and acetonitrile with 0.1% trifluoroacetic acid were used as the mobile phase. The analytical response, represented as relative peak area ratios (analyte to internal standard), was converted to amounts extracted by employing an instrumental calibration curve consisting of analytes in the desorption solvent in the 0.1- 100 ng/mL range.

### Safety analysis

Adult rats were administered either saline vehicle (n=3) or ketamine-loaded liposomes (n=3) via IV infusion. Ultrasound was applied to all animals after 2.5 minutes of drug infusion for 2.5 minutes (250 kHz, 0.9 MPa estimated in situ peak negative pressure, 50 ms pulse length, 5 Hz PRF, 25% duty cycle). The targeted site was 5 mm posterior to bregma and +2.5 mm lateral to midline to the right side of the brain. At 72 hours post-infusion, rats were euthanized and fixed via transcardial perfusion with 1x PBS and 4% paraformaldehyde (PFA). Brains were extracted and stored in 4% PFA for 24 hours, cryopreserved in 15% sucrose for 48 hours, and lastly, in 30% sucrose for 48 hours. Brains were frozen in optimal cutting medium compound and stored at -80 °C until serially sectioned with a cryostat at 30 μm in the horizontal plane. Every 12th section (360 μm apart) was stained with hematoxylin and eosin (H&E), Fluorojade-C, GFAP, and IBA-1 to evaluate for parenchymal damage, neuronal degeneration, and astrocytic or microglial activation, respectively. Alexa Fluor® 488 secondary antibody was used after primary anti-GFAP incubation. Cy5® secondary antibody was used after primary anti-IBA1 incubation. Tissue sections were free-float mounted on microscope glass slides (Fisher, Pittsburgh, PA). All histology images were collected with a fluorescence microscope (BZ-X800, Keyence Corp., Itasca, IL). For quantifiable histological markers (Fluoro-Jade C, IBA-1+, and GFAP+ cells), signals above thresholded background were used for manual ROI segmentation to calculate the total mean fluorescent area of cells using BZ-X Advanced Analysis Software (Keyence Corp., Itasca, IL).

### Von Frey Experiment

Rats were acclimated to a raised stainless steel mesh table for a minimum of 15 minutes. They were then anesthetized and kept under 2% isoflurane during the experiment. Saline vehicle or ropivacaine-loaded liposomes was administered, followed by focused ultrasound applied using a dorsal approach to target the sciatic nerve. Paw withdrawal response was evaluated using monofilaments and the Von Frey up-down method 30 minutes, 1 hour, and 4 hours post treatment, as previously described (*63, 64*).

### Statistical analysis

GraphPad Prism 5 (GraphPad Software, La Jolla, CA) was used for statistical analysis. Comparisons between two groups were performed by two-tailed Student’s t-test, and that among multiple groups by one way analysis of variance (ANOVA) with Dunnett’s post hoc multiple comparisons tests. A p-value of 0.05 or less was considered statistically different and the p-values were categorized as *: p < 0.05, **: p < 0.01, and ***: p < 0.001.

## List of Supplementary Materials

Fig. S1: Norketamine & hydroxynorketamine biodistribution with ketamine-loaded AAL versus unencapsulated ketamine-HCl infusion.

Fig. S2: Quantification of brain histological markers to assess for safety with ultrasonic ketamine uncaging.

## Acknowledgements

We would like to thank Kim Butts Pauly, Jeremy Dahl, Katherine Ferrara, Sami Karaborni, Vivianne Tawfik, Boris Heifets, Andrew Neice, Jeffrey B. Wang, and the whole Airan Lab for helpful discussions. We would also like to thank the Stanford CryoEM center (cEMc) including Dr. Elizabeth Montabana and Dr. Bharti Singal for technical assistance in cryo-EM imaging.

## Author Contributions

Formulated AALs and designed in vitro analyses (MPP, YX, RDA); Performed in vitro analyses (MPP, YX, ARH, DGL); Designed and calibrated HPLC, LC/MS-MS, and SPME protocols and performed SPME/LC/MS-MS analysis (KSR); Designed in vivo analyses (KSR, YX, BJY, GM, RDA); Performed in vivo analysis and sample acquisition (KSR, YX, BJY, GM, AKT, DGL); Performed histologic procedures and analysis (MMA); Performed data analysis (MPP, KSR, YX, BJY, MMA, GM, ARH, RDA); Drafted the figures and manuscript (MPP, KSR, YX, BJY, MMA, AKT, RDA); Funding acquisition (RDA).

## Data and materials availability

Data and code are available from the corresponding author upon reasonable request.

