## Supplemental Figures for "Acoustomechanically activatable liposomes for ultrasonic drug uncaging"

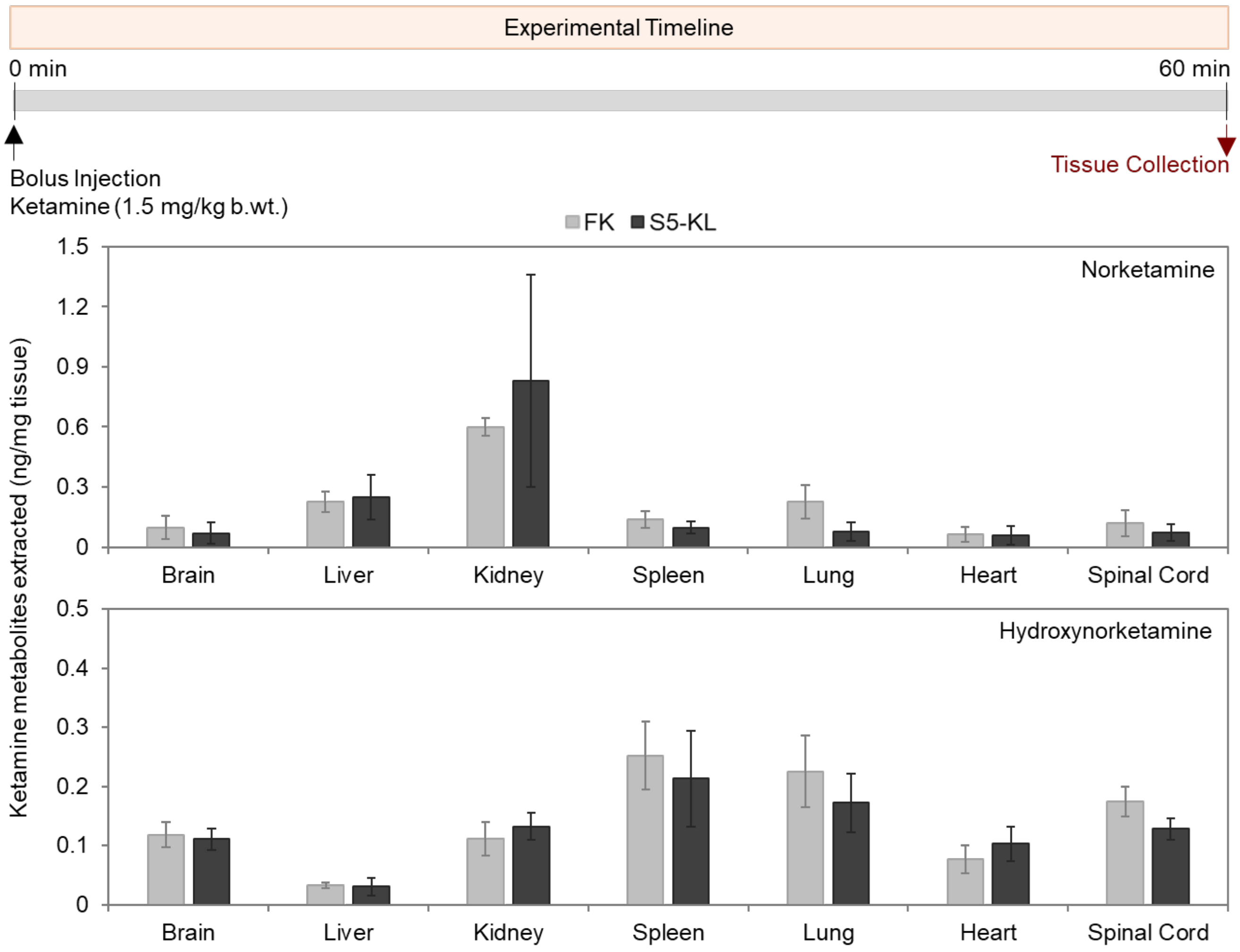


**Fig S1.** Norketamine & hydroxynorketamine amounts extracted from each organ of the biodistribution experiment reported in Fig. 4.

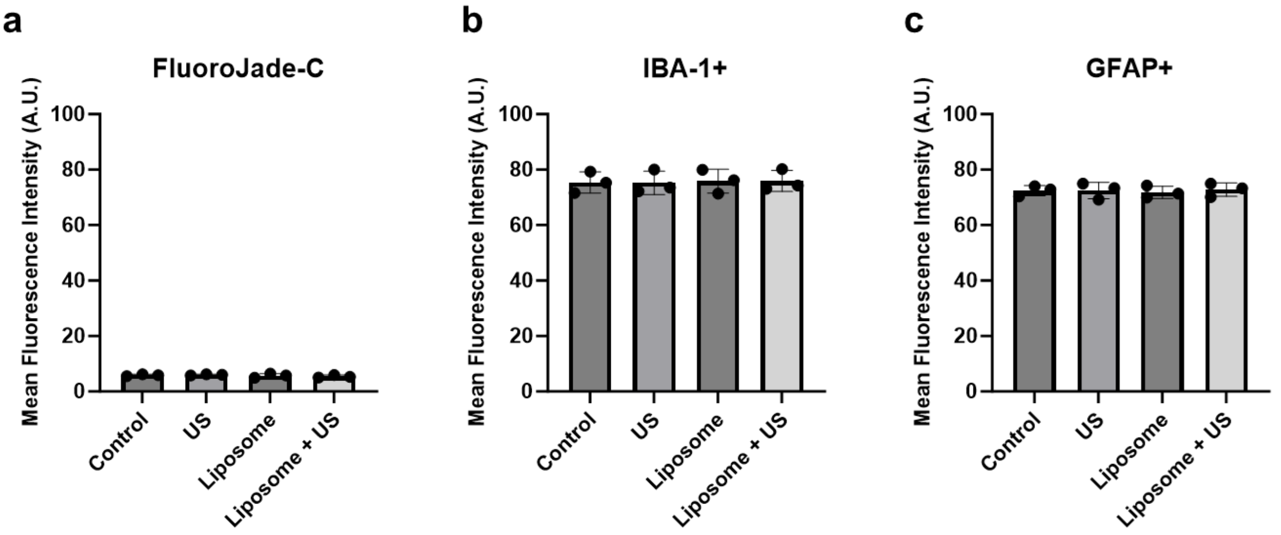


**Fig S2**. Quantification of brain histological markers to assess for safety following ultrasonic ketamine uncaging. **(a)** Mean fluorescence intensity of total FluoroJade C positive cells (marker of neuronal degeneration). In comparison to healthy control brains, there was no difference in levels of FluoroJade C positive cells in either Ultrasound (US), liposome, or liposome+US groups. **(b)** Mean fluorescence intensity of total IBA-1 positive cells (marker of microglial activation). In comparison to healthy control brains, there was no difference in levels of IBA-1 positive cells in either US, liposome, or liposome+US groups. **(c)** Mean fluorescence intensity of total GFAP positive cells (marker of astrocytic activation and gliosis). In comparison to healthy control brains, there was no difference in levels of GFAP positive cells in either US, liposome, or liposome+US groups. Data presented as mean ± S.D., analyzed by one-way ANOVA with post-hoc Dunnett's multiple comparisons test; N=3 for each group.
